# Sex-biased gene regulation varies across human populations as a result of adaptive evolution

**DOI:** 10.1101/2023.04.12.536645

**Authors:** Adam Z. Reynolds, Sara D. Niedbalski

## Abstract

Human males and females exhibit a wide range of diversity in biology and behavior. However, studies of sexual dimorphism and gender disparities in health generally emphasize ostensibly universal molecular sex differences, such as sex chromosomes and circulating hormone levels, while ignoring the extraordinary diversity in biology, behavior, and culture acquired by different human populations over their unique evolutionary histories. Using RNA-Seq data and whole genome sequences from 11 human populations, we investigate variation in sex-biased gene expression across human populations and test whether population-level variation in sex-biased expression may have resulted from adaptive evolution in sex-specific regulatory regions. In tests for differential expression, we find that sex-biased gene expression in humans is highly variable, mostly population-specific, and demonstrates between-population reversals. Expression quantitative trait locus (eQTL) mapping reveals sex-specific regulatory regions with evidence of recent positive natural selection, suggesting that variation in sex-biased expression may have evolved as an adaptive response to ancestral environments experienced by human populations. These results indicate that sex-biased gene expression is more flexible than previously thought and is not generally shared among human populations. Instead, molecular phenotypes associated with sex depend on complex interactions between population-specific molecular evolution and physiological responses to contemporary socioecologies.

## INTRODUCTION

Humans exhibit sexually dimorphic characteristics and biological processes across virtually all body systems (Dunsworth, 2020; Frayer & Wolpoff, 1985; Klein & Flanagan, 2016; Naqvi et al., 2019; Stulp & Barrett, 2016). Gender disparities in health and disease are also widespread (Agyemang et al., 2008; Swinburn et al., 2011). Further, human males and females display an extraordinary range of diversity across populations, in both behavior and biology (Low, 2015; Roughgarden, 2013; Ubelaker & DeGaglia, 2017). However, studies of sexual dimorphism in humans generally focus on the causes and consequences of ostensibly universal molecular differences between males and females (e.g., sex chromosomes, steroid hormones), while ignoring variation acquired by different populations over the course of their unique evolutionary histories (Austad, 2006; Frayer & Wolpoff, 1985; Klein & Flanagan, 2016; Rigby & Kulathinal, 2015). As ancestral humans dispersed across the globe, they entered—and adapted to—a variety of novel environments (Moreno-Mayar et al., 2018; Nielsen et al., 2017; Pagani et al., 2016), encountering many opportunities for evolution to impact sexually dimorphic traits. However, there remains little clarity on the extent to which human males and females vary molecularly across populations or whether adaptive evolution over the course of human history has contributed to meaningful biological variation in sexual dimorphism.

Here we investigate the landscape of human variation in sex-biased gene expression and assess whether global patterns of sex-biased expression have resulted from adaptive evolution in genomic regulatory sequences. Sex-biased expression occurs in thousands of genes that are distributed across the autosomal chromosomes as well as the sex chromosomes (Dimas et al., 2012; Ellegren & Parsch, 2007; Melé et al., 2015), and varies by tissue, developmental stage, and disease context (Arnold et al., 2009; Mayne et al., 2016; Naqvi et al., 2019). While sex-linked genes on the X or Y chromosome do play a role in sex-specific patterns of gene expression, the complexity of *cis-* and *trans-*regulatory networks and the scale of age- and tissue-specific signatures of sex-biases in autosomal genes make it unlikely that the majority of sex-biased expression is driven by the sex chromosomes (Grath & Parsch, 2016; Melé et al., 2015; Naqvi et al., 2019; Oliva et al., 2020). Similarly, although there are many genes that harbor androgen or estrogen response elements in their promoters or in known enhancer regions, only about 32% of sex-biased genes are regulated by sex hormones (Mayne et al., 2016; Shen et al., 2017). Thus, sex-linked genes and circulating sex hormones do not generally account for the wide range of sexually dimorphic molecular phenotypes in humans.

Studies of long-term evolution reveal that sex-biased genes evolve more rapidly than unbiased genes, both in sequence and regulation (Assis et al., 2012; Ellegren & Parsch, 2007; Grath & Parsch, 2016; Meiklejohn et al., 2003; Naqvi et al., 2019; Ranz et al., 2003). While sex biases in gene expression are modestly conserved across the evolutionary lineages of mammals, most appear to have been acquired recently and the particular bias observed in any given gene can vary substantially across tissues and species (Ellegren & Parsch, 2007; Mayne et al., 2016; Naqvi et al., 2019; Rigby & Kulathinal, 2015). Patterns of sex-biased expression seen in humans are generally not shared with other animals (Naqvi et al., 2019) and, specifically, signals of selection in male-biased genes have been discovered on the human lineage following divergence from chimpanzees (Nielsen et al., 2005). Evidence of recent adaptation has been discovered in unbiased genes contributing a range of phenotypes, including immune responses (Nédélec et al., 2016; Sanz et al., 2018), height (Lopez et al., 2019; Perry et al., 2014), vitamin D metabolism (Amorim et al., 2017; Hlusko et al., 2018), and high-altitude adaptation (Brutsaert et al., 2019; Buroker et al., 2012), as well as specifically in *cis*-regulatory expression quantitative loci (eQTLs) (Kudaravalli et al., 2009; Romero et al., 2012). Although hundreds of *cis-*eQTLs that regulate sex-biased genes have been identified in humans (Mayne et al., 2016; Naqvi et al., 2019; Oliva et al., 2020), these studies have been limited to single populations, so they have been unable to test whether the observed sex biases are universal across human populations.

Here we investigate the hypothesis that sex biases in gene regulation and expression vary across human populations and test whether population-specific differences in sex-biased expression may have resulted from adaptive evolution in sex-specific *cis*-regulatory variants. Using data from 11 populations with diverse evolutionary histories and living in a variety of contemporary environments, we demonstrate that sex biases in gene expression are not generally shared among human populations. We then conduct sex-specific eQTL mapping and tests for natural selection to assess whether sex-specific gene regulation may be the result of population-specific adaptation over the course of human dispersal. Signatures of positive selection in *cis-*regulatory variants indicate that variation in sex-biased expression may have evolved as an adaptive response to the differing historical environments experienced by human populations.

## MATERIALS AND METHODS

### Data Sources and Preparation

All data used in this study are publicly available as part of the Thousand Genomes (1000G) and the Human Genome Diversity (HGDP) projects. RNA sequence data were collected by Lappalainen et al. (Lappalainen et al., 2013) and Martin et al. (Martin et al., 2014) from EBV-transformed lymphoblastoid cell lines (LCLs), which reliably preserve a snapshot of the mRNA at the time of cell capture and transformation (Çalışkan et al., 2011). Matched whole genome sequences come from the 1000G phase 3 (The 1000 Genomes Project Consortium, 2015) and HGDP (Bergström et al., 2019) sequencing efforts. We excluded two samples from the 1000G due to poor read quality as well as all samples from the San population (HGDP), because they included RNA sequences only from males. We also excluded two Cambodian males, one Yakut male, and one Yakut female because they did not have matching WGS available in HGDP. Our final dataset was comprised of 465 samples from 1000G (95 Finnish, 94 British, 91 Utah, 93 Toscani, 89 Yoruba) and 41 samples from HGDP (7 Cambodian, 7 Maya, 7 Mbuti, 7 Mozabite, 7 Pathan, 6 Yakut). Figure 1 shows the populations included in the analysis. Figure 2 outlines our analysis strategy.

**Figure 1.**
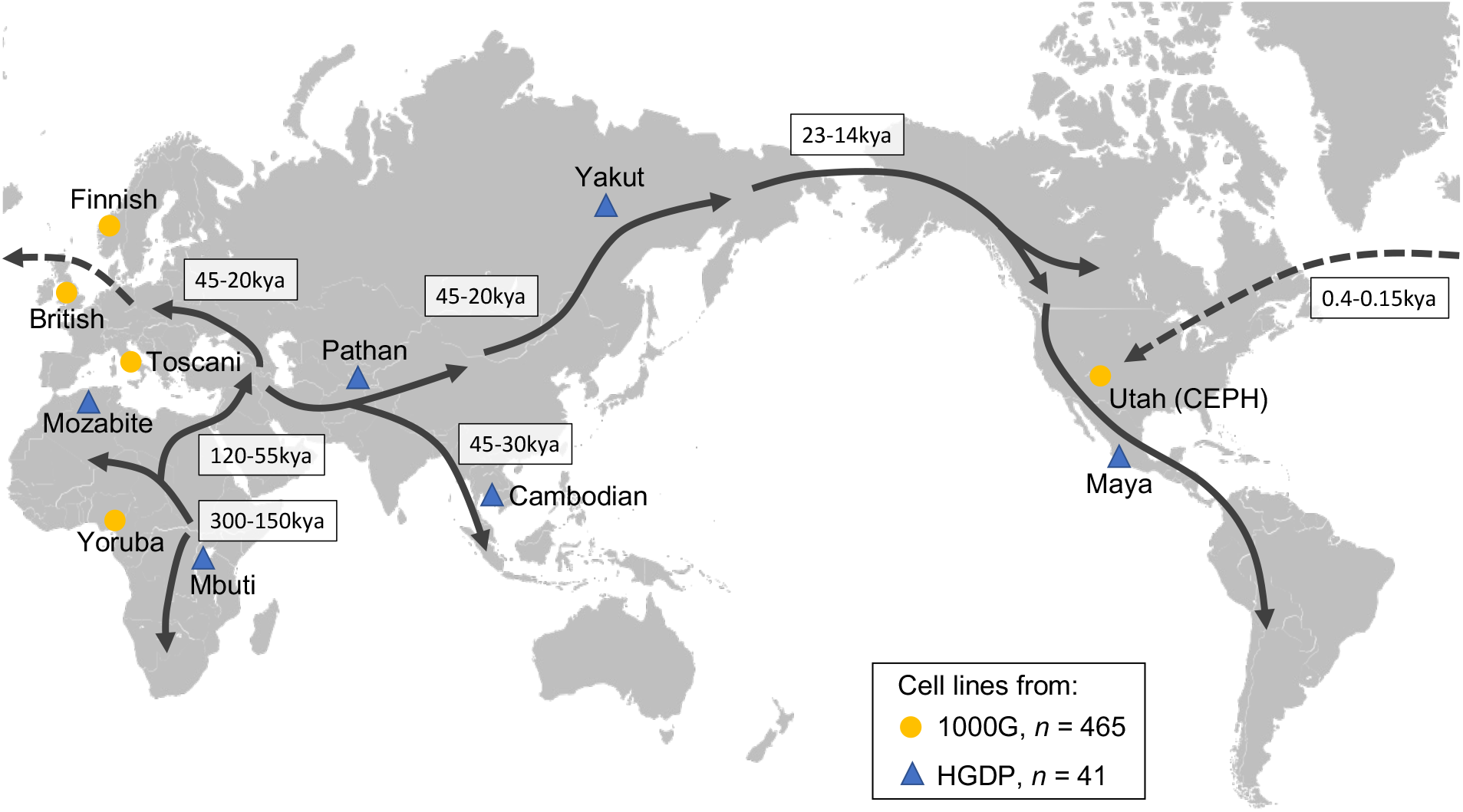
Populations included in the analysis. Arrows depict the history of human dispersal across the globe (Bergström et al., 2021; Choin et al., 2021; Dausset et al., 1990; Pagani et al., 2016; Raghavan et al., 2015; Schiffels & Durbin, 2014). Human genetic variation is patterned by serial founder effects, which reduced genetic diversity from Africa to the Americas (Ramachandran et al., 2005), as well major population bottlenecks, which allowed novel alleles to go to high frequencies during migrations out-of-Africa, into Oceania, and into the Americas (Niedbalski & Long, 2022). Selection for sex-biased gene expression may have occurred in ancestral environments over the course of human dispersal or in recent environments experienced since population divergence.

**Figure 2.**
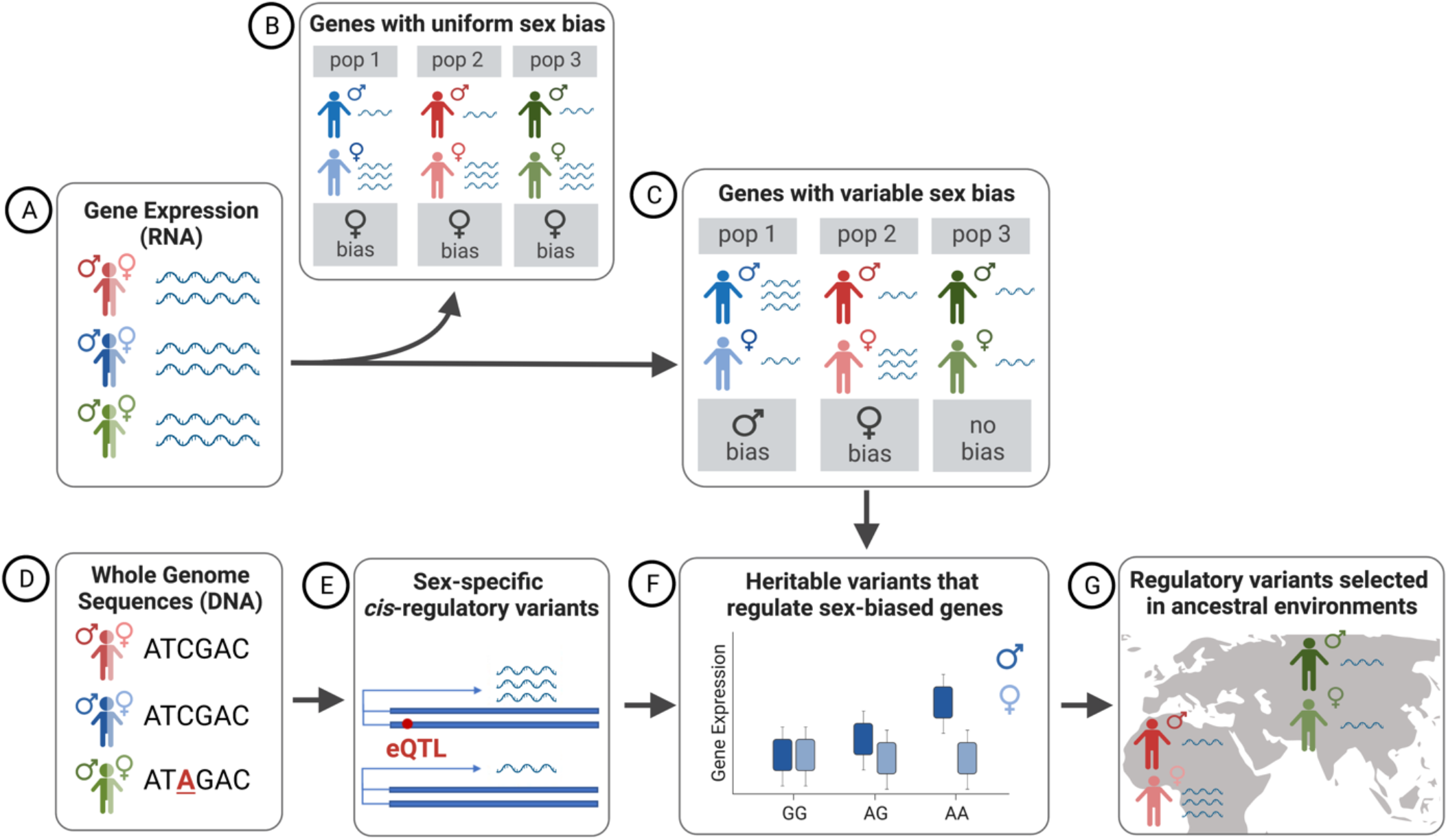
Analytical strategy. Using RNA-Seq data (A), we conduct differential expression analysis with a sex*population interaction to distinguish genes exhibiting a signal of universal sex bias (B) from genes exhibiting variable sex bias across populations (C). With matched whole genomes (D), we then identify sex-specific *cis*-regulatory variants (E) that explain variability in sex-biased genes (F). Finally, we test whether heritable genetic variants that regulate sex-biased genes appear to have undergone natural selection in ancestral environments (G).

### Differential Gene Expression Analysis

Prior to analysis, we aligned paired-end RNA-Seq reads to the human reference genome version GRCh38/hg19 using *HISAT2* (D. Kim et al., 2019, p. 2) and then counted uniquely-aligning reads using *featureCounts* version 1.6.4 (Liao et al., 2014). To identify sex-biased gene expression within and across populations, we tested for differential expression between males and females using the *DESeq2* package (Love et al., 2014) in R version 4.1.0 (Core Team, 2020). *DESeq2* uses a generalized linear model with dispersion shrinkage to estimate the presence and strength of differential expression based on a matrix of read counts. All models were fitted using beta priors and sample-specific size factors. Tests of significance were conducted using a negative binomial Wald Test. To account for multiple tests, p-values were adjusted using a Benjamini-Hochberg correction with a false discovery rate (FDR) of 0.1.

Due to differences in sample size and sequencing platforms between the 1000G and HGDP RNA-Seq data, it was not feasible to adequately control for batch effects in a single analysis. Thus, differential expression analysis was conducted separately on each dataset. To assess whether biased expression in particular genes varies across populations, we modeled an interaction between sex and population, using design equation *∼sex*population*. To determine whether gene expression is sex-biased in each particular population, we tested for the main effect of sex in the reference population and report genes for which this term had a significance of *p* < 0.05 after FDR correction. To identify whether some genes exhibit variation in sex bias across populations, we tested whether the FDR-corrected term for the interaction of sex and population was statistically significant at the same threshold.

### Expression Quantitative Trait Locus (eQTL) Mapping

To assess whether sex-biased gene expression might be a product of gene regulatory variants, we tested for *cis-*expression quantitative trait loci (*cis-*eQTLs), defined here as regions containing one or more expression-associated single nucleotide polymorphisms (SNPs) located within one megabase of the transcription start site. To do this, we matched our normalized gene read count tables with whole genome sequence data for the same individuals from 1000G and HGDP. eQTL analysis of 1000G samples was conducted in human genome build 37, while samples from HGDP were analyzed in build 38. SNPs with a minor allele frequency (MAF) > 0.05 and located within a 1 Mb region of transcription start site for each gene were considered. To test for association of each SNP with gene expression levels, we used a linear model, which assumes an additive effect of the number of copies of each variant. Significant associations were identified using a p-value threshold of 1e-3 following a Benjamini-Hochberg correction with FDR=0.1. All models were conducted using the *matrixEQTL* package (Shabalin, 2012) in R (Core Team, 2020).

### Tests for Positive Natural Selection

To test for evidence of adaptive evolution in sex-biased eQTLs, we employed *Relate* (Speidel et al., 2019), which uses a coalescent-aware modeling approach to estimate allele frequency trajectories and calculate *p*-values on the strength of selection throughout population history. To do this, *Relate* uses a hidden Markov model and knowledge of the ancestral and derived alleles at each site to estimate the relative timing of historical coalescence events between a particular haplotype and all other haplotypes, then constructs a rooted binary tree with branch lengths estimated using a Bayesian Markov chain monte carlo approximation that allows for changes in effective population size. Relate estimates a *p*-value for selection by comparing the speed of spread of a mutation in a particular lineage relative to other ‘competing’ lineages present when the mutation first arises. In our tests for selection, we included WGS data for all available individuals in the populations of interest from the 1000G and HGDP datasets. Prior to analysis, all WGS were recoded to use ancestral/derived allele status and phased using the scripts provided with *Relate* software. Each site was determined to exhibit evidence of positive selection if calculated p-values were < 0.05.

## RESULTS

### Variability in sex-biased expression across diverse populations

We find that sex-biased gene expression is highly variable and largely population-specific. Differential expression analysis revealed 22 genes with evidence of shared sex bias across populations. Importantly, we detected zero genes with universal bias across all eleven populations studied. Instead, these 22 genes were detected in all 5 populations of 1000G, but not in the lower-power HGDP dataset. Except for two autosomal genes, *LINC01597* (chr20) and *EIF2S3B* (chr12), which are universally female- and male-biased, respectively, the remainder were female-biased and located on chrX. When we broaden our definition to include a more moderate signature of universal bias—genes with the same sex bias in at least two populations and no significant sex*population interaction—we find an additional 14 genes on the X chromosome, two of which (*ARMCX1* and *OCRL*) are male-biased in expression, as well as an additional 23 genes in the autosomal chromosomes, of which 10 are female-biased and 13 are male-biased. Among these, three female-biased ribosomal protein genes (*RPS4XP3, RPS4XP6*, and *RPS4XP7*) were detected in four populations, while the male-biased *ZPBP2* was detected in three populations. Table 1 lists genes showing at least a moderate signature of universal sex bias.

**Table 1.**
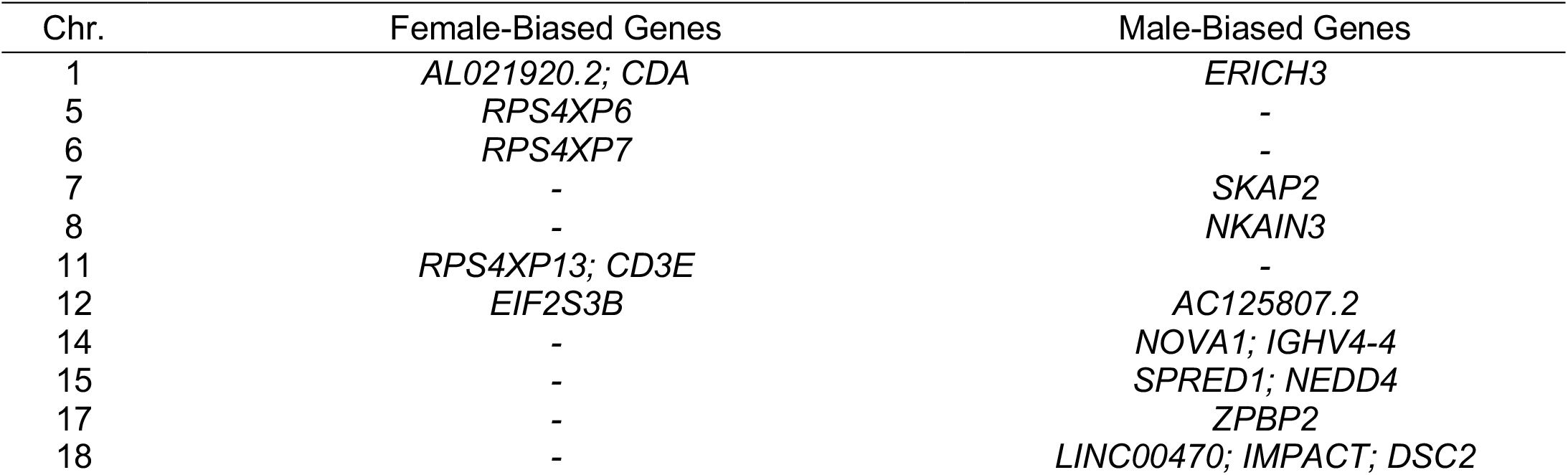
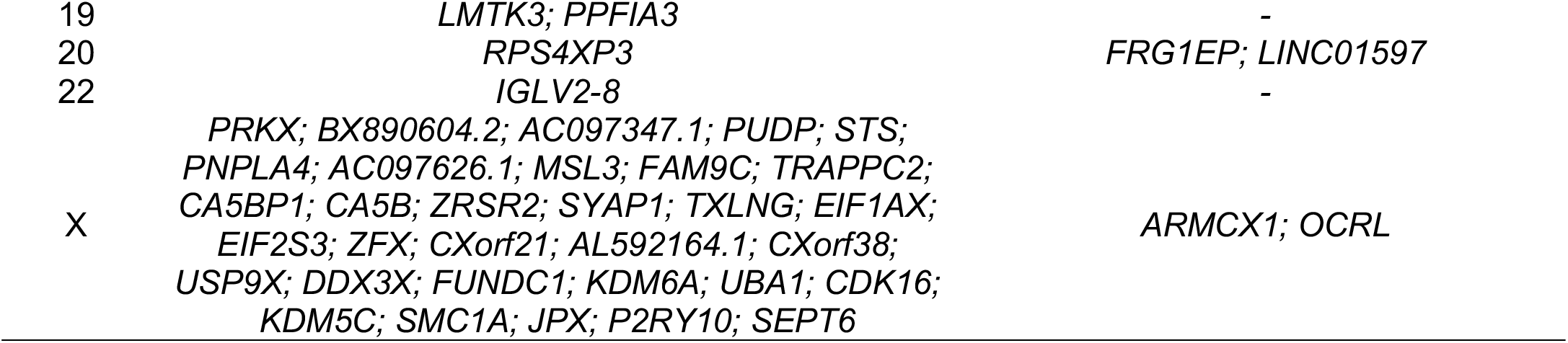
Genes with moderate evidence of universal sex bias.

More generally, populations in our analysis show diversity in the number, log fold change, and particular genes exhibiting sex-biased expression. The distributions in Figure 3 reveal a wide range of variation in the relative counts and strengths (log2 fold change) of sex bias for genes with male-biased (negative) and female-biased (positive) expression across populations. Among 1000G populations, for example, the Yoruba exhibit many more genes with significantly sex-biased expression than we observe in the European populations. Moreover, although the distributions for Utah and the Yoruba are relatively symmetrical, this is not the modal pattern among populations in our analysis. Most groups exhibit greater counts of genes with male-biased expression, while the Cambodians are the only population to exhibit an overall female bias. However, in the Utah, Maya, and Pathan populations, the strongest sex biases are observed in genes with female-biased expression.

**Figure 3.**
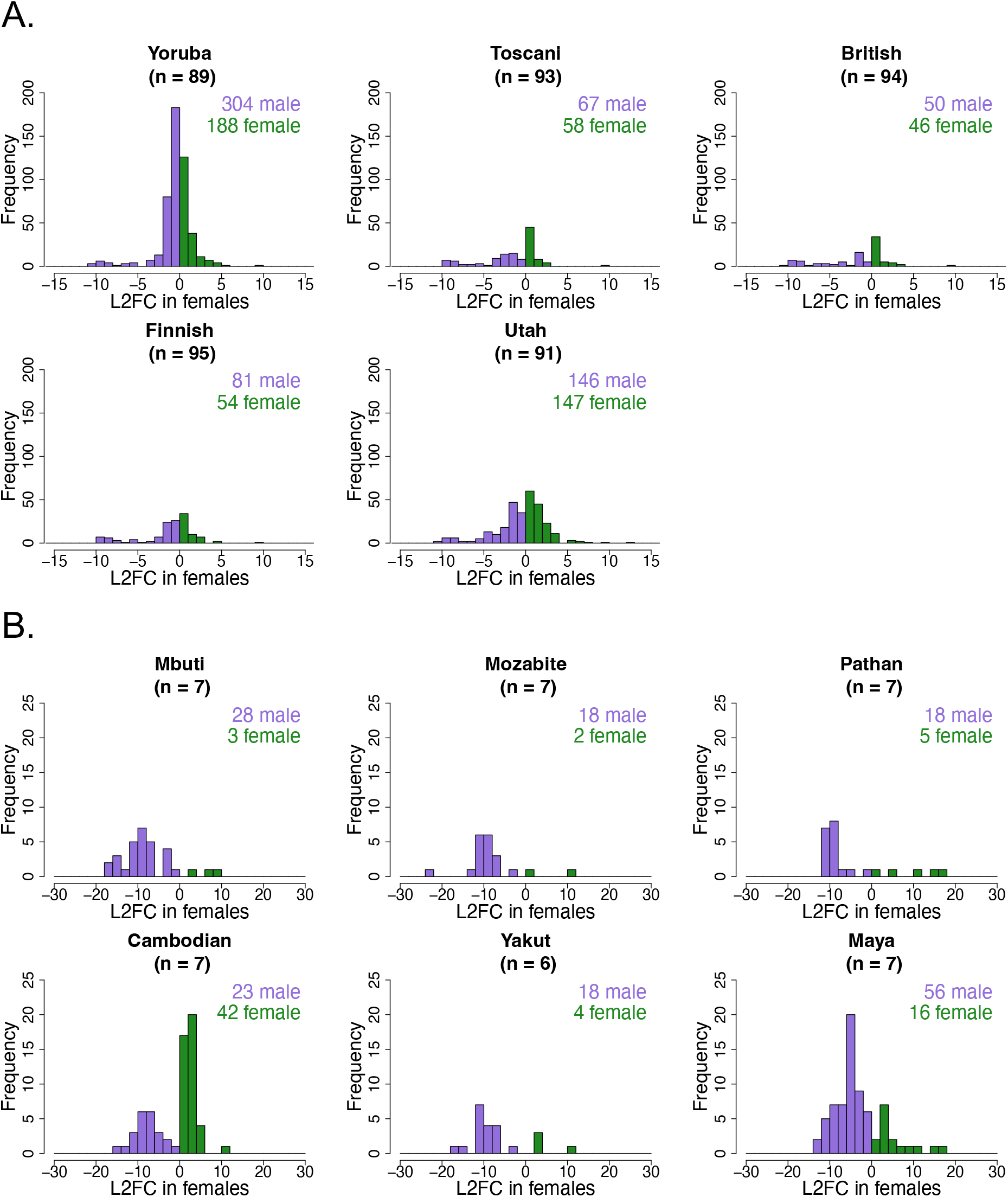
Distributions of genes with sex-biased expression in populations from the 1000G (A) and HGDP (B). Log2 fold change (L2FC) indicates the strength of sex-biased expression. Genes with positive L2FC are female-biased; genes with negative L2FC are male-biased.

We detected FDR-corrected significant sex*population interactions for 278 unique genes in the 1000G dataset and 76 genes in the HGDP data, yielding a total of 345 unique genes for which sex bias varies significantly across human populations. As shown in Figure 4, the large majority of these were found to be sex-biased in only a single population, with the highest counts observed in Utah (118) and the Yoruba (90). These two populations also combined for the highest number of shared genes (5), while only one other pair had more than one (2 were shared between Utah and the Maya). Overall, very few sex biases were found to be shared among other pairs or triplets of populations.

**Figure 4.**
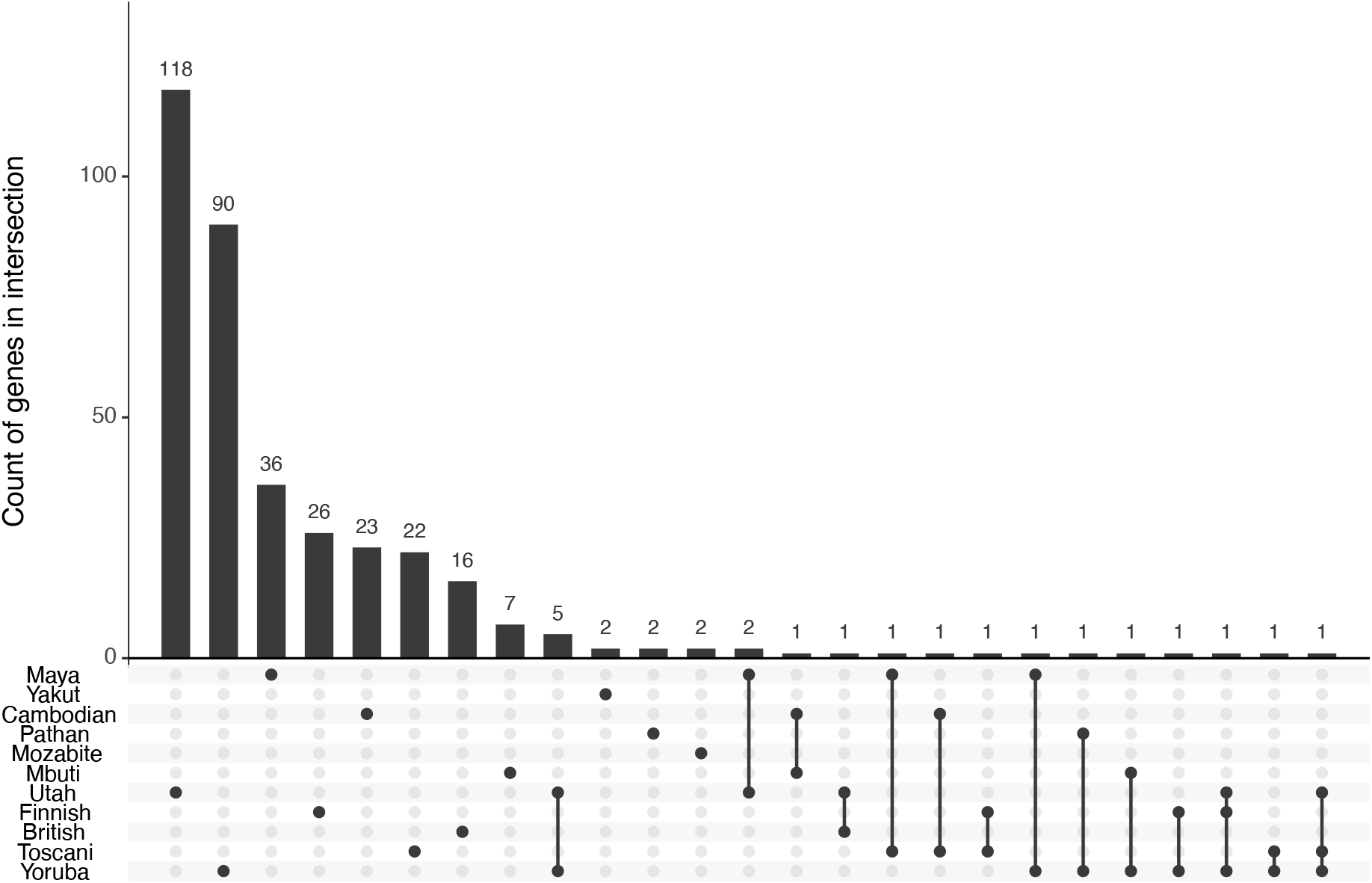
Genes with shared sex bias across populations. Plot contains only genes for which there was a significant interaction of sex*population; genes exhibiting universal sex bias are reported in text and in Table 1 above.

### Cross-population reversals in sex-biased expression

Intriguingly, our analyses reveal 27 genes that exhibit reversed sex bias across at least one pair of populations (Table 2). As seen in Figure 5, these reversals are distributed widely across the genome and vary between different pairs of populations. Helped by the larger sample sizes, most reversals were detected in the 1000G populations, but we were able to detect reversals in genes with relatively strong bias among the HGDP populations. Most of these involve reversals between one of Yoruba or Utah and the other European populations, while a few also reverse between the Mbuti and Maya. In light of the population-specific nature of sex-biased expression and the variety of populations involved in at least one reversal, it is difficult to tease apart the possible contributions of ancestry and environment to reversals in sex-biased gene expression.

**Table 2.**
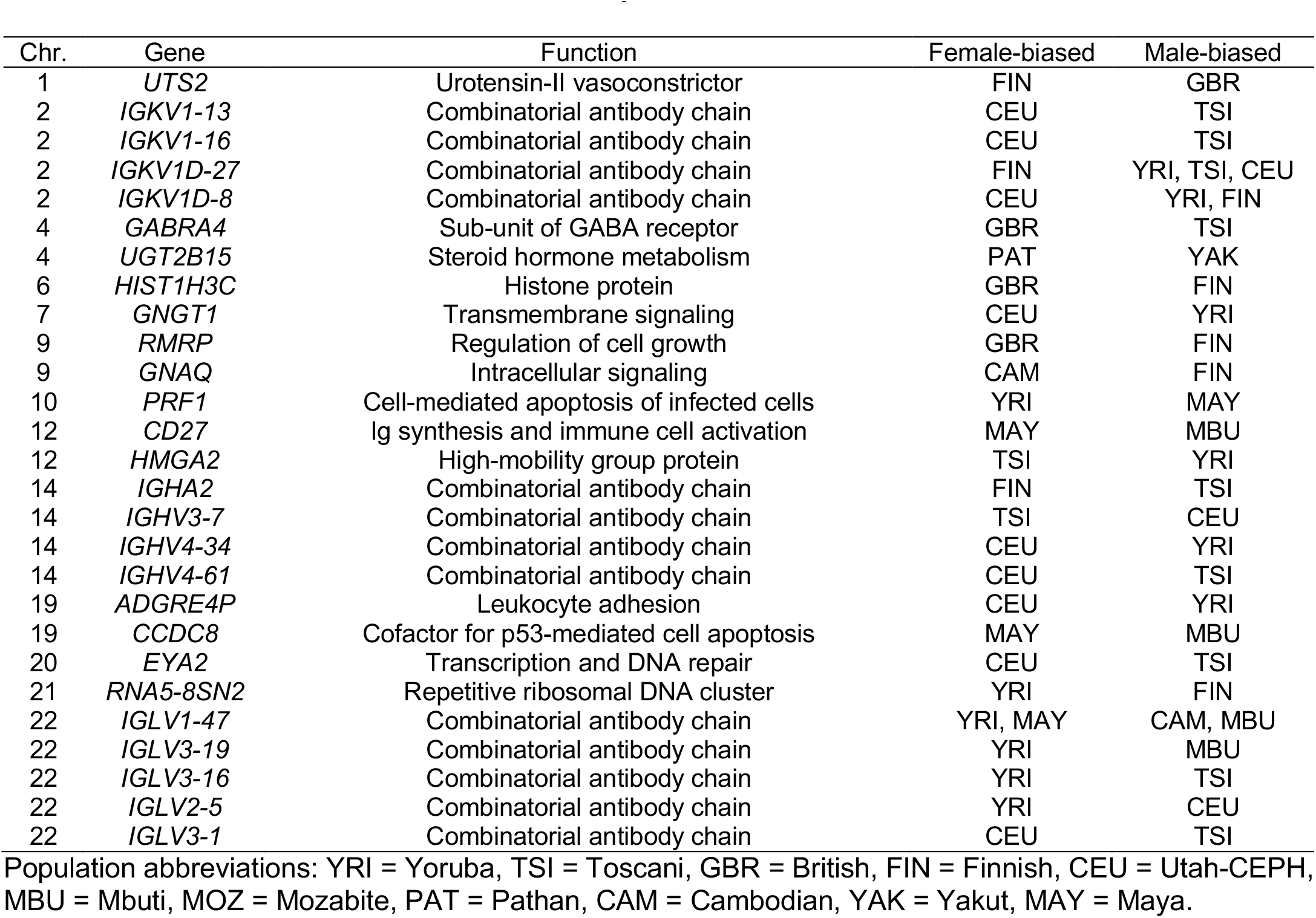
Genes with reversals in sex-biased expression.

**Figure 5.**
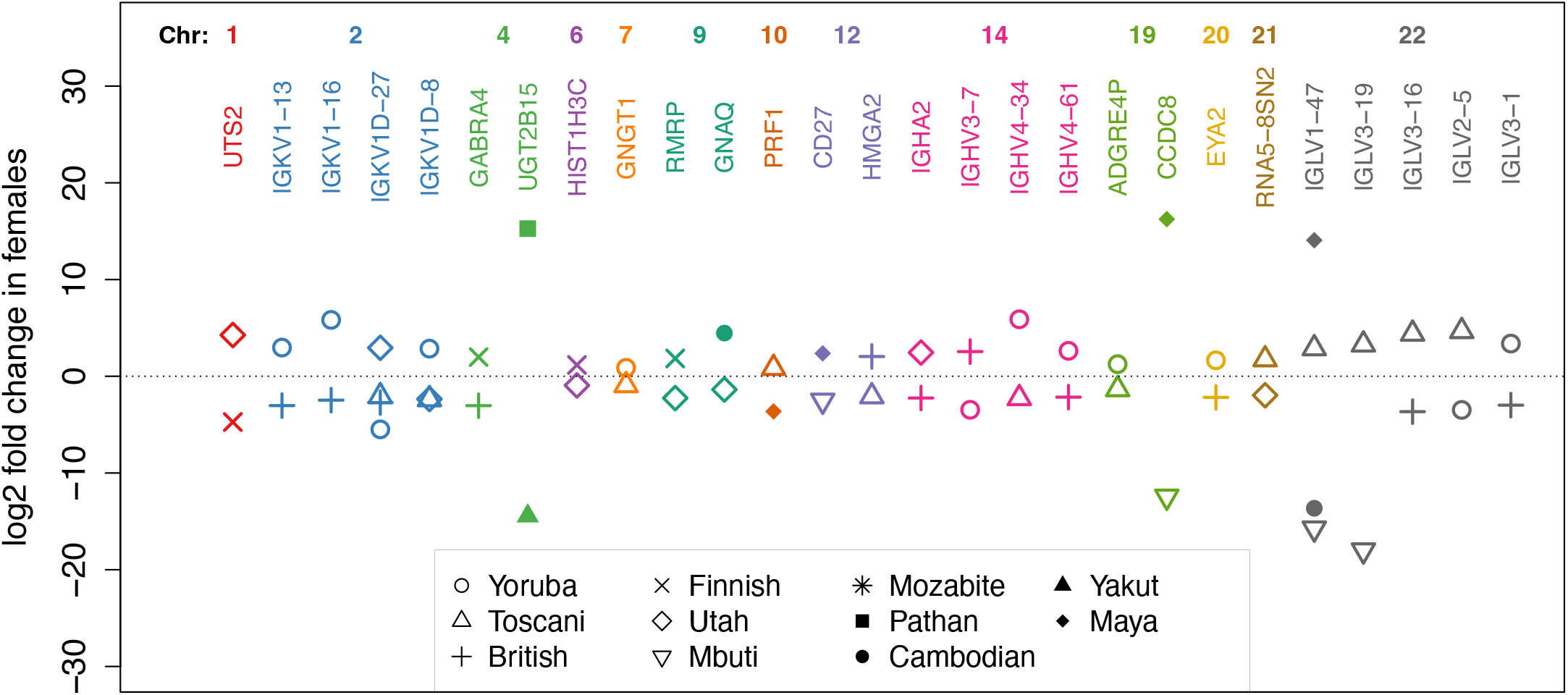
Twenty-seven genes exhibit reversed sex bias across populations. Log2 fold change indicates how many times a gene’s expression is doubled in females compared to males. Negative values indicate male bias, while positive values indicate female bias. Populations with unbiased expression not shown.

Thirteen genes with reversing sex-biased expression code for various chains in the immunoglobulin (Ig) complex, indicating that males and females are expressing different combinations of antibodies in different populations. In the 1000G dataset, reversals in the Ig complex are apparent among the Utah-CEPH, Toscani, Yoruba, and, to a lesser extent, the Finnish. In contrast, the British do not exhibit sex biases in any of these genes. In the HGDP dataset, we observe a reversal in the light variable chain *IGLV1-47*, which is female-biased in the Maya (and Yoruba) and male-biased in both Mbuti and Cambodians. We also find reversals in two other genes participating in adaptive immunity: *CD27* (female-biased in the Maya, male-biased in the Mbuti), which up-regulates Ig synthesis and activates differentiation of memory B-cells into plasma cells (Agematsu et al., 2000; Buchan et al., 2018), and *GABRA4* (female-biased in the British, male-biased in the Toscani), a sub-unit of the GABA receptor with an anti-inflammatory role (J. K. Kim et al., 2023; Yocum et al., 2017). Given that our data were sequenced from lymphoblastoids, it is perhaps unsurprising that we detect significant variation in genes participating in immune function. However, the observation of reversals in adaptive immune genes highlights that some sex biases in gene regulation may be facultative responses to variation in socioecology, such as sex-specific pathogen exposures in particular populations.

Reversals are also apparent in genes affecting a number of other systems. *UGT2B15*, which codes for a catalyst of sex-steroid hormone metabolism, is female-biased in the Pathan and male-biased in the Yakut, indicating that even processes regulating levels of circulating sex hormones may reverse between populations. We also observe that urotensin-II (*UTS2*), the strongest known vasoconstrictor with important roles in cardiac, nervous, and renal system function, is female-biased in the Finnish and male-biased in the British. Additionally, we find reversals in a number of genes participating in cell cycle regulation (*RMRP, HMGA2, CCDC8, EYA2*), cell signaling (*GNAQ, PRF1, GNGT1*), and DNA structural regulation (*HIST1H3C, HMGA2*). A number of these genes have been linked to chronic diseases and cancers (Jiang et al., 2008; Markowski et al., 2013; Pang et al., 2018; Pereira-Castro et al., 2019; Revelo et al., 2014; Wang et al., 2017; Zhang et al., 2019), which raises the possibility that reversals in sex-biased gene expression may put males and females in different populations at differing risk for chronic diseases or adverse outcomes from these diseases.

### Natural selection on eQTLs regulating sex-biased expression

To identify whether variation in sex-biased expression is heritable and may have resulted from adaptive evolution, we conducted expression quantitative trait locus (eQTL) mapping and then tested for evidence of natural selection in genomic regions containing sex-specific eQTLs. A substantial number of genes with variable sex bias are influenced by heritable regulatory variants: eQTLs regulating female expression were detected in 121 sex-biased genes, while another 127 genes had eQTLs that regulate male expression. Tests for selection revealed 33 regions in 1000G and 2 additional regions in HGDP that exhibit evidence of selection on regulatory variants contributing to sex-biased expression (Table 3). Genes showing signatures of selection are widely distributed across the autosomal chromosomes. Lists of SNPs and chromosomal regions are available in Supplemental Table S1.

**Table 3.** Selected eQTLs regulating sex-biased gene expression.

| Chromosome | Gene | Selected SNPs | Population<sup>a</sup> | Expression Bias<sup>b</sup> | Effect of Selected SNPs<sup>c</sup> |
| --- | --- | --- | --- | --- | --- |
| 1 | *UTS2* | 3 | GBR | M | ↓ F |
| 1 | *ZMYM4* | 6 | YRI | M | ↓ F |
|  |  | 26 | EUR | U | ↑ F |
| 2 | *LINC01115* | 1 | YRI | M | ↑ M |
| 2 | *TFPI* | 3 | FIN | M | ↑ M |
| 3 | *CAND2* | 4 | CEU | F | ↓ M |
| 3 | *ADAMTS9* | 13 | YRI | M | ↑ M |
|  |  | 2 | EUR | U | ↓ M |
| 4 | *TENM3* | 2 | YRI | M | ↑ M |
|  |  | 2 | EUR | U | ↓ M |
| 6 | *SERPINB1* | 2 | MAY | M | ↑ M |
| 6 | *PNLDC1* | 14 | YRI | F | ↑ F |
|  |  | 7 | TSI, GBR, CEU | U | ↓ F |
| 6 | *HIST1H1C* | 7 | YRI | F | ↑ F |
|  |  | 11 | EUR | U | ↓ F |
| 6 | *MAP3K5* | 2 | YRI | M | ↑ M |
| 7 | *MEST* | 2 | TSI | M | ↑ M |
| 7 | *DPP6* | 4 | YRI | M | ↑ M |
|  |  | 1 | TSI, GBR | U | ↓ M |
| 8 | *MSR1* | 1 | YRI | M | ↑ M |
|  |  | 1 | TSI, FIN, CEU | U | ↓ M |
| 8 | *RIMS2* | 6 | YRI | M | ↑ M |
|  |  | 9 | EUR | U | ↓ M |
| 9 | *AIF1L* | 3 | YRI | M | ↑ M |
|  |  | 2 | TSI, GBR, CEU | U | ↓ M |
| 10 | *PRF1* | 2 | YRI | F | ↑ F |
| 10 | *CTBP2* | 6 | YRI | M | ↑ M |
|  |  | 2 | TSI, GBR, FIN | U | ↓ M |
| 11 | *ARHGAP32* | 1 | CEU | M | ↓ F |
| 12 | *CD4* | 1 | CEU | M | ↑ M |
| 12 | *TMEM119* | 11 | YRI | F | ↑ F |
|  |  | 1 | TSI, CEU | U | ↑ M |
| 13 | *RNASEH2B-AS1* | 6 | FIN | F | ↓ M |
| 13 | *EDNRB* | 1 | TSI | M | ↑ M |
| 14 | *PTGER2* | 2 | CEU | M | ↓ F |
| 14 | *SERPINA9* | 1 | YRI | M | ↑ M |
|  |  | 2 | EUR | U | ↓ M |
| 14 | *SERPINA1* | 2 | TSI | M | ↑ M |
| 14 | *IGHV3-65* | 1 | YRI | F | ↑ F |
|  |  | 2 | TSI, GBR, FIN | U | ↑ M |
| 14 | *IGHV4-34* | 1 | YRI | M | ↑ M |
| 18 | *EPB41L3* | 1 | CEU | M | ↑ M |
| 18 | *CBLN2* | 11 | YRI | M | ↑ M |
| 19 | *JSRP1* | 11 | YRI | F | ↑ F |
|  |  | 3 | EUR | U | ↓ F |
| 19 | *CREB3L3* | 19 | YRI | F | ↑ F |
|  |  | 1 | EUR | U | ↓ F |
| 22 | *CECR2* | 3 | YRI | M | ↑ M |
|  |  | 1 | EUR | U | ↓ M |
| 22 | *IGLV1-47* | 1 | MBU | M | ↑ M |
| 22 | *IGLV2-5* | 2 | YRI | F | ↑ F |

<sup>a</sup> Population: YRI = Yoruba, TSI = Toscani, GBR = British, FIN = Finnish, CEU = Utah, EUR = all four populations of European descent (TSI, GBR, FIN, CEU), MBU = Mbuti, MAY = Mayan.
<sup>b</sup> Sex bias: F = female, M = male, U = unbiased.
<sup>c</sup> Regulatory effect of selected SNPs: ↑M = up-regulating male expression, ↓M = down-regulating male expression, ↑F = up-regulating female expression, ↓F = down-regulating female expression.

Figure 6A shows genes with selected regulatory regions in the 1000G, shaded by the strength of evidence for selection in each population. While 23 of these show selection on eQTLs contributing to male-biased expression, just 10 eQTLs show evidence of selection for female-biased expression. Several genes show evidence of selection in just a single population suggesting they result from relatively recent selective events on a single lineage following divergence from the other populations. In 22 genes, we detected evidence of selection favoring sex-biased expression in the Yoruba, far more than were discovered in the Toscani (3), British (1), Finnish (2), and Utah (5) combined. However, we also observed that 16 of these carry a distinct signature of two-way selection both *for* sex-biased expression in the Yoruba as well as *against* sex-biased expression in European populations. Representative allele frequency trajectories for the *ZMYM4* gene (Figure 6B) show that SNPs contributing to reduced sex bias in Europeans began increasing in frequency around 50kya. This suggests that there may have been important differences between the historical environments of Yorubans (favoring sex-biased expression) and ancestral Europeans (favoring unbiased expression) during or after the out-of-Africa migration, but before divergence of European populations. Genes regulated by these regions are involved in a range of systems and processes, including the inflammatory response (*CREB3L3*), bone remodeling and ossification (*TMEM119*), regulation of cholesterol storage (*MSR1*), white fat cell differentiation (*CTBP2*), skeletal muscle contraction (*JSRP*), and morphogenesis of the aorta, arteries, and cardiac muscle tissues (*ADAMTS9*).

**Figure 6.**
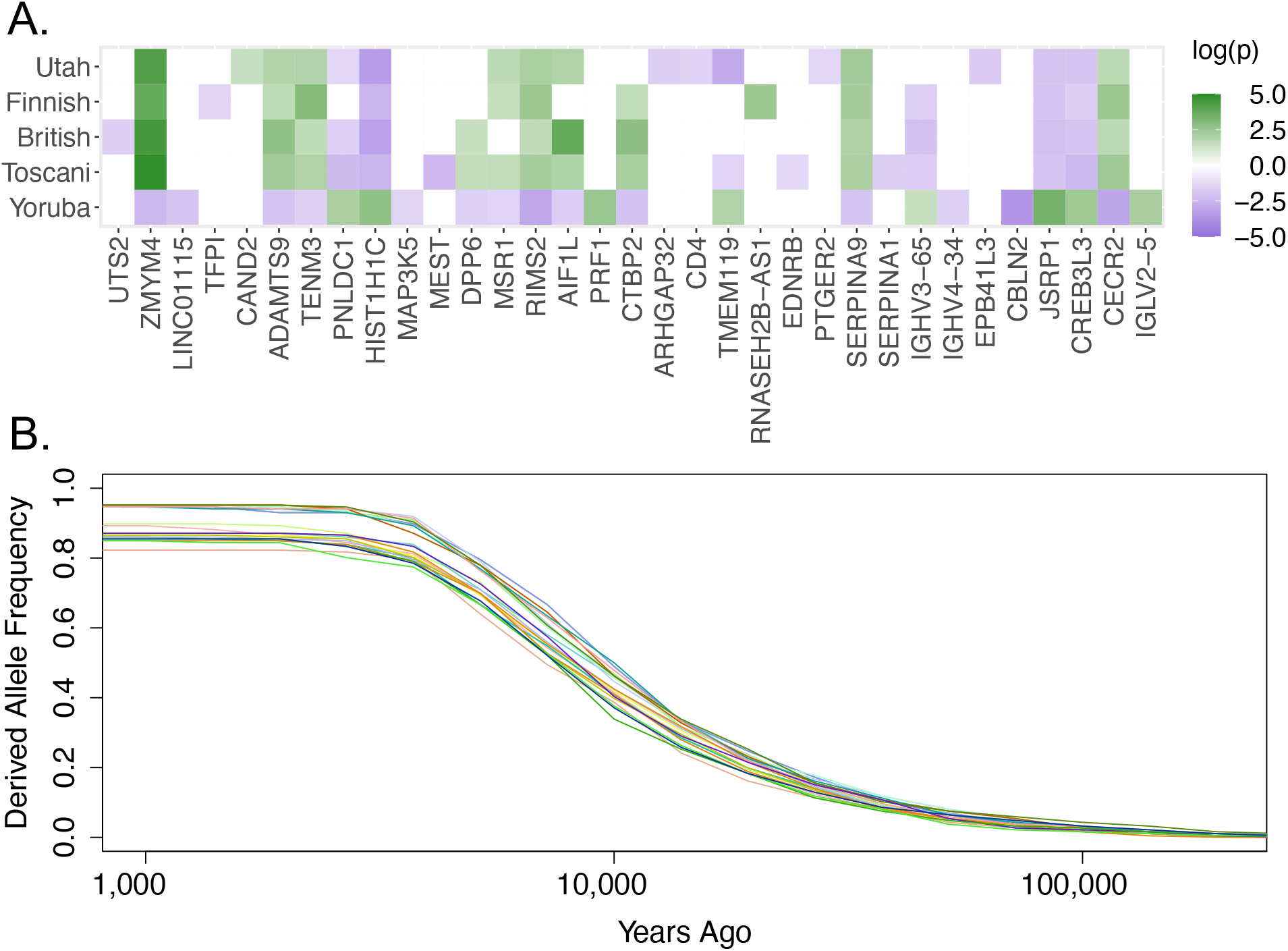
Evidence of selection on eQTLs regulating sex-biased genes. (A) Genes with evidence of selection on regulatory eQTLs in the 1000G populations. Sixteen genes show a two-way signal of selection, favoring both sex bias in the Yoruba and reduced sex bias in Europeans. Selection in the direction of female-biased expression (or away from male-biased expression) is represented by positive log10 p-values (green), while selection in the direction of male-biased expression (or away from female-biased expression) is represented by negative log10 p-values (purple). (B) Representative allele frequency trajectory for selected variants contributing to reduced sex bias in Europeans (*ZMYM4* shown). Coalescent-estimated historical allele frequencies suggest selection for reduced sex bias beginning ∼50kya, after the out-of-Africa migration but before divergence of European populations.

We also find that natural selection may be responsible for some genes reversing in sex bias across populations. Specifically, we detect evidence of selection in sex-specific eQTLs regulating *UTS2, PRF1, IGLV2-5*, and *IGLV1-47*. Notably, all four of these genes participate in highly facultative responses to environmental factors: The Ig complex (*IGLV1-47, IGLV2-5*) forms antibodies that provide immunity against previously encountered pathogens, perforin (*PRF1*) is an immune protein critical for the destruction of infected cells, and urotensin-II (*UTS2*) regulates cardiovascular, renal, and neurological responses to environmental stressors. However, while we observe reversed sex biases in these genes across populations, we detect signals of selection for only one directional effect in each case. For example, while we find evidence of selection on a region that down-regulates female expression of *UTS2* in the British (thereby contributing to male bias), we do not find a similar signal on any variants that could contribute to the female bias we observe in the Finnish. Similarly, in *IGLV2-5*, we detect evidence of selection on variants contributing to female bias in the Yoruba, but no comparable signal on variants contributing to the observed male bias in Utah. These results highlight that population-level variation (and reversals) in sex-biased gene expression are due in part to the combined processes of evolution by natural selection and shorter-term physiological responses to the environment.

## DISCUSSION

As humans dispersed across the globe, they encountered—and adapted to—a range of novel environments, producing extraordinary diversity in human biology, behavior, and culture. In this study, we investigated the extent to which sexual dimorphism in gene expression is shared across human populations and find, by contrast, that sex-biased gene expression is both highly variable and mostly population-specific. Previous studies have demonstrated that sex-biased expression varies broadly across tissues and is not primarily regulated by sex-linked factors (Mayne et al., 2016). In the present study, we demonstrate that genes exhibiting sex bias in one population are not generally the same genes that exhibit sex bias in other populations. Indeed, the regulation of sex-biased gene expression in a Toscani male differs substantially from that of a Yoruban, Mayan, or Finnish male. Thus, previously-reported signals of sex bias detected in samples from a single population (Mayne et al., 2016; Melé et al., 2015; Naqvi et al., 2019) are unlikely to generalize to other human groups.

In fact, sex-biased gene expression varies to such a degree that some genes even completely reverse in sex bias across populations. While it has previously been shown that sex biases in expression can reverse across tissues (Mayne et al., 2016; Melé et al., 2015; Oliva et al., 2020), we here demonstrate that this is also true across populations, even within the same cell type. Thus, the human genome cannot be neatly classified into male-biased and female-biased genes: “male genes” in one context may in fact be “female genes” in another context. Instead, the genomic regulatory features underlying sexually dimorphic molecular phenotypes appears to be much more facultative and context-dependent than portrayed in previous studies. In general, gene expression is widely variable and highly responsive to behavioral and environmental factors, including diet, sleep, cold stress, and urban environments (Aho et al., 2013; Cole, 2014; Idaghdour et al., 2008, 2010; Kelly et al., 2017; Kohrt et al., 2016; Ornish et al., 2008; Shinozaki et al., 2003). Many of the sex-biased genes identified in the present study are known to respond facultatively to the presence of particular environmental stimuli. For example, some of the most flexible genes we observe are involved in humoral immunity, including a number of Immunoglobulin (Ig) chains that participate in combinatorial antigen recognition. Reversals in these genes may result from socioecological variation across human populations that differentially exposes males and females to particular pathogens.

Our results demonstrate that variation in sex-biased gene expression results in part from molecular adaptation to the ancestral environments experienced by human populations over their unique evolutionary histories. Although environmental variation almost certainly influences patterns of sex-biased gene expression across human populations, these sex biases can, over evolutionary time, become encoded in the genome via positive natural selection on sex-specific regulatory variants. Our results thus concord with other studies arguing that sex-biased genes can evolve relatively rapidly (Grath & Parsch, 2016; Ranz et al., 2003). However, whereas previous studies have emphasized evolution between species, the present analysis demonstrates that there has been enough time within the history of the human species for evolution on heritable molecular variation contributing to population-specific patterns of sex-biased gene regulation.

In nearly half of the genes for which we observed a selected *cis*-regulatory eQTL, we detected a two-way pattern of evolution both maintaining sex-biased expression in Yorubans and reducing sex-biased expression in Europeans. Reduced sexual dimorphism in Europeans has been hypothesized to be a result of more similar types of labor performed by men and women in agricultural societies (Frayer, 1980). Our results, however, align with other studies refuting this hypothesis (Arner et al., 2021; Holden & Mace, 1999; Wolfe & Patrick Gray, 1982): Although we detect a signal of selection on eQTLs contributing to reduced sex bias in Europeans, it appears the selective event would have occurred during or after the out-of-Africa migration, but prior to divergence between European groups, thus pre-dating the onset of agriculture.

An alternative hypothesis with more empirical support suggests that sexual dimorphism in body size is related to the amounts, rather than types, of physical activity undertaken by males and females (Buffa et al., 2001; Holden & Mace, 1999). Consistent with this hypothesis, our observation of a relatively early selective sweep for reduced sex bias in Europe coincides with archaeological evidence for both increased physical activity and limited sexual dimorphism among European hunter-gatherers during the Upper Paleolithic (Villotte et al., 2010). Importantly, the presence of these signals in all four populations of European descent and the estimated timing of their allele frequency increases leaves open the possibility that they did not result from natural selection at all, but in fact could have drifted to high frequency during the extreme population bottleneck following the out-of-Africa migration (Nei et al., 1975; Rogers & Mukherjee, 1992).

Taken together, our analyses uncover wide variation in sex-biased gene expression among human populations with unique evolutionary histories and living in diverse environments. These observations are relevant for understanding risk factors that affect sex and gender disparities in health and disease. The overwhelming majority of studies on sex differences in health have been conducted in western, industrialized, and patriarchal societies, leading to hypotheses that sex/gender disparities are driven by universal biological differences between males and females (for reviews, see (Klein & Flanagan, 2016; Natri et al., 2019). However, a growing body of evidence from traditionally under-represented populations suggests that in fact sex and gender disparities in many diseases are more variable than previously believed (Abarca-Gómez et al., 2017; Mills et al., 2020; Reynolds et al., 2020; Zhou et al., 2017). The results of the present study imply that our understanding of sex-linked risk factors and sex as a biological variable (Clayton, 2018; Stachenfeld & Mazure, 2022) will increase in complexity as genomic epidemiology studies incorporate more diverse populations. Sex and gender disparities in health are not due solely to universal biological differences between males and females, nor is variation in these disparities solely a product of cultural and behavioral diversity; instead, the impact of sex on health and disease depends on complex interactions between population-level genetic adaptation to ancestral environments and physiological responses to contemporary socio-ecologies.

## ACKNOWLEDGEMENTS

Training support for this work was provided by a New Mexico IDeA Network for Biomedical Research Excellence (NM-INBRE) Award by the National Institute of General Medical Sciences (5P20GM103451). Computing resources and technical assistance were provided by the staff at the National Center for Genome Resources (NCGR) in Santa Fe, NM

